# Pep2Mol: 3D Molecule Generation Targeting Protein-Protein Interfaces with Diffusion Models

**DOI:** 10.64898/2026.06.28.734975

**Authors:** Rongting Yue, Zekun Yang, Gustavo Seabra, Chenglong Li, Yanjun Li

## Abstract

Protein-protein interactions (PPIs) are central to biological processes. Designing small molecules that modulate dysregulated PPIs holds strong promise for targeting undruggable proteins. However, existing structure-based drug design approaches focus on well-defined small-molecule binding pockets and struggle to generalize to large, shallow, and chemically complex PPI interfaces. Here, we introduce Pep2Mol, a diffusion-based generative model for 3D molecule design that targets orthosteric PPI sites by explicitly incorporating binding peptides or proteins as structural guidance, moving beyond conventional pocket-conditioned generation. To enable model development and benchmarking, we curate a large-scale, high-quality dataset of 10,956 experimentally resolved protein complex structure pairs, each pairing an orthosteric competitive ligand with a protein binder at overlapping receptor interfaces. Pep2Mol integrates two SE(3)-equivariant graph neural networks that encode protein–ligand and protein–peptide interactions respectively, and fuses these representations via attention-based conditioning to jointly guide the diffusion trajectory. Extensive evaluations demonstrate that Pep2Mol generates chemically valid ligands with state-of-the-art binding affinities, providing a strong foundation for small-molecule inhibitor design against challenging PPI interfaces.

## 1 Introduction

Protein-protein interactions (PPIs) play a fundamental role in biological processes such as signal transduction, cell proliferation, differentiation, and apoptosis Lu et al. [2020], Greenblatt et al. [2024]. The human interactome is estimated to contain 74, 000 to 200, 000 PPIs Venkatesan et al. [2009]; dysregulation of these interactions is implicated in a wide range of diseases. Although small molecules offer a promising drug discovery strategy for modulating PPIs by targeting their interfaces, their development remains highly challenging Nero et al. [2014], Shin et al. [2020].

Structure-based drug design (SBDD) exploits three-dimensional protein structures to guide the rational design of small molecules that bind specific pockets with high affinity Batool et al. [2019]. Recent advances in deep generative models have substantially accelerated this paradigm, enabling the automated generation of ligands tailored to defined binding regions Tang et al. [2024]. However, most existing approaches have been developed and evaluated primarily on canonical ligand-binding pockets, which are typically compact, well-enclosed, and enriched in hydrophobic residues. By contrast, PPI interfaces exhibit fundamentally different biochemical and geometric characteristics, being generally larger, flatter, more flexible, and dominated by extended surface contacts Wang et al. [2024], Ni et al. [2019], Shin et al. [2017]. Owing to these differences, current approaches often demonstrate suboptimal performance when applied to PPI interfaces.

In addition, atomic interactions between target proteins and protein or peptide binders at PPI interfaces encode valuable structural information that is often leveraged by medicinal chemists to guide small-molecule inhibitor design. For example, peptidomimetic strategies aim to develop small molecules that mimic the binding modes and key interactions of native peptide binders at protein-protein interfaces Pelay-Gimeno et al. [2015]. However, few existing generative models effectively exploit the rich interaction information embedded in PPI structural data, substantially limiting their capacity.

To address these limitations, we introduce Pep2Mol, a full-atom, diffusion-based generative framework for 3D molecule design that targets orthosteric PPI interfaces by explicitly incorporating binding peptides or proteins as structural guidance. By unifying the PPI and protein-ligand interaction (PLI) contexts within a single framework, our model effectively captures and leverages atomic-level interaction information derived from peptide binders during the diffusion process, thereby guiding the design of orthosteric small-molecule inhibitors with strong target binding affinity.

The development of this framework requires a large, structurally rich dataset that links PPIs to their corresponding ligand-binding structures. To this end, we curate a high-quality dataset of experimentally resolved PPI-PLI pairs from the Protein Data Bank (PDB), capturing orthosteric binding scenarios in which small-molecule inhibitors competitively bind at protein-protein interaction interfaces. Unlike existing collections such as 2p2idb Basse et al. [2016], which provide limited protein family-level coverage, our curated dataset focuses on pocket-level data and encompasses 10, 956 high-quality pairs forming, to the best of our knowledge, the largest dataset available dedicated to PPI interfaces with known orthosteric modulators. This extensive dataset provides a solid foundation for learning the fine-grained geometric and physicochemical cues that govern orthosteric PPI modulation. Building on our curated dataset, Pep2Mol employs a diffusion-based framework, in which the forward diffusion gradually perturbs the ligand coordinates and atom types. During reverse diffusion, the protein interface (or “hot spots”) serves as the initial structural condition, and the model learns to denoise a randomly initialized ligand. Protein or peptide binder information is integrated via a cross-attention mechanism and equivariant message passing using SE(3)-equivariant graph neural networks (EGNNs) Satorras et al. [2021]. Pep2Mol effectively fuses key interaction features of PPI interfaces, preserving geometric consistency while enriching the context for ligand generation. The overview of our model is illustrated in Figure 1.

**Figure 1:**
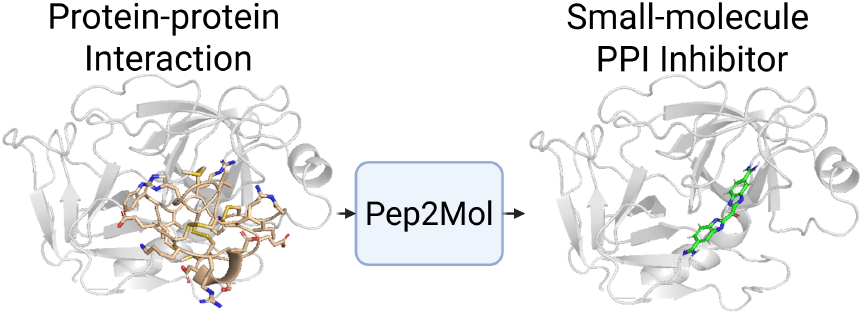
Illustration of Pep2Mol. Starting from a PPI complex, Pep2Mol generates 3D small-molecule inhibitors that bind the interaction interface by distilling the key interaction patterns of the protein binder into a compact small-molecule form.

In summary, our contributions are: (1) we introduce Pep2Mol, a diffusion-based SBDD framework for the de novo generation of 3D orthosteric small-molecule inhibitors targeting PPI interfaces, explicitly guided by the protein-protein interaction information beyond typical pocket-conditioned generation. (2) we curate the largest high-quality, machine learning-ready dataset to date of 10, 956 experimentally resolved orthosteric PPI–PLI pairs, providing a new foundational resource for structure-based PPI inhibitor design and AI-driven drug discovery; (3) extensive experimental evaluations demonstrate that Pep2Mol generates high-affinity ligands targeting challenging PPI interfaces.

## 2 Related Work

### 2.1 Structure-based Drug Design

Structure-based drug design exploits 3D protein structures to enable the rational design of small molecules. Early generative approaches focused on producing 1D SMILES strings or 2D molecular graphs conditioned on protein contexts Skalic et al. [2019], Tan et al. [2023], Wang et al. [2025]. Subsequent methods advanced toward direct 3D generation, employing variational autoencoders and autoregressive models to produce ligand conformations explicitly aligned with receptor binding sites Liu et al. [2022], Peng et al. [2022], Ragoza et al. [2022], Zhang et al. [2023]. For example, Luo et al. [2021] propose a 3D AR model that samples atom coordinates conditioned on protein binding sites. Pocket2Mol Peng et al. [2022] adopts an autoregressive atom- and bond-placement strategy guided by local pocket geometry. Denoising diffusion probabilistic models Ho et al. [2020] have recently become a more commonly used strategy for 3D structure-based ligand generation Guan et al. [2023a,b], Francoeur et al. [2020], Huang et al. [2024a], Zheng et al. [2026]. In this setting, diffusion models denoise molecular representations conditioned on protein environments. For example, TargetDiff Guan et al. [2023a] introduces a pocket-conditioned diffusion model based on an SE(3)-equivariant network, enabling effective pocket-aware ligand generation. IRDiff Huang et al. [2024a] augments diffusion with retrieved high-affinity ligand references from a pretrained protein– ligand interaction network, while IPDiff Huang et al. [2024b] incorporates interaction priors into both the forward and reverse diffusion processes, achieving improved affinity and structural realism on CrossDocked2020 Francoeur et al. [2020]. More recently, Apo2Mol Zheng et al. [2026] further accounts for pocket dynamics and enables ligand generation from unbound pocket states. Besides the diffusion models, DrugFlow Shen et al. [2024] employs a flow matching-based framework for generating ligand atom type and coordinates and bond types conditioned on the target pocket. Despite their promise in SBDD, these approaches primarily focus on canonical ligand-binding pockets and remain limited in their ability to generalize to large, shallow, and chemically complex PPI interfaces.

### 2.2 Discovery of PPI Inhibitors

PPIs represent therapeutically important yet challenging drug targets, motivating substantial efforts to develop computer-guided strategies for identifying small-molecule PPI inhibitors. Existing methods, however, have primarily focused on predicting inhibitory potency or distinguishing PPI inhibitors from non-inhibitory compounds Krishnan et al. [2021], Wolk and Goldblum [2022], Gao et al. [2023], Zhang et al. [2024]. Generative modeling for PPI inhibitors remains largely underexplored. iPPI-GAN Wang et al. [2022] applies a GAN-based framework with voxelized molecular representations and 3D CNNs to capture shape complementarity at PPI interfaces, followed by SMILES decoding. GENiPPI Wang et al. [2024] explores PPI-aware generation by conditioning on protein complex interface features, but is evaluated on only a small set of PPI targets and does not explicitly constrain ligand generation to the binding site. Unlike the standard ligand-binding pockets, in PPI targets, protein binders define precise orthosteric interfaces and provide atomic-level interaction patterns, including key contact residues and binding motifs that can be mimicked by competitive small molecules. Therefore, incorporating protein binder information provides a promising and biologically grounded conditioning signal for structure-based generative design of orthosteric PPI inhibitors.

## 3 Preliminary

### 3.1 Problem Statement

In this work, we investigate orthosteric PPI inhibitor generation, where the goal is to generate a small-molecule ligand that competitively binds to a PPI interface defined by a peptide binder. In our dataset, each data instance consists of a receptor protein, a peptide or protein binder, and a corresponding small-molecule ligand that binds to the orthosteric PPI interface. For simplicity, throughout this paper, we use the term “peptide” to refer to peptide binders and “PPI interface” to refer to interface segments extracted from receptor proteins in PPI complexes, consistent with our dataset construction. However, for PPI targets to which our framework is applicable, the binders are not restricted to short peptides and can also include full-length proteins.

The ligand-binding pocket of the receptor protein is represented as 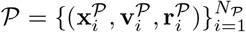, where 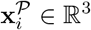 denotes atomic coordinates, 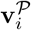denotes atom-type features, and 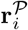 denotes residue-type features. The aligned peptide is represented as 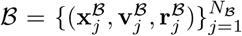, where 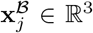 denotes atomic coordinates, 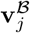 denotes atom-type features, and 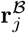denotes residue-type features of the peptide atoms. The binder *B* is represented in the same aligned coordinate frame as the receptor pocket *P* and provides peptide-derived structural guidance for competitive ligand generation. The target ligand is denoted as 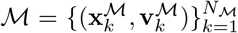, where 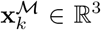 and 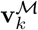 are the ligand atom coordinates and atom-type features, respectively, and *N*_*M*_ is the number of atoms.

The learning objective is to model the conditional distribution *p*(*M*| *P, B*), i.e., generating a small-molecule ligand conditioned jointly on the protein pocket and the aligned peptide binder. Unlike conventional pocket-based ligand generation, the PPI interface provides explicit structural cues about orthosteric binding patterns, guiding the design of competitive PPI inhibitors.

### 3.2 Diffusion Models in SBDD

*M*

*P*

Diffusion models Ho et al. [2020], Song et al. [2021] are a class of generative models that have achieved notable success across diverse domains. They define a forward process that progressively perturbs data through noise injection and learn a reverse denoising process for sample generation. Recently, diffusion models have been widely adopted for SBDD tasks Guan et al. [2023a], Huang et al. [2024b], Schneuing et al. [2024], Zheng et al. [2026]. In this setting, the protein pocket and ligand molecule *M* are typically represented by atomic coordinates **x** ∈ ℝ^3^ and atom-type features **v**, and the molecule generation is formulated as modeling the conditional distribution *p*(*M* | *P*). At each diffusion step *t* in the forward process, the ligand state, including continuous atomic coordinates 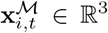 and discrete atom types 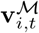, is corrupted by a predefined noise schedule until *M*_*T*_ approaches a Gaussian prior:

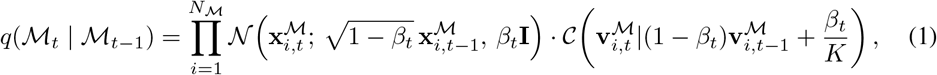

where *N* and *C* represent Gaussian and categorical distributions, respectively, *K* is the number of atom-type categories, and the noise schedules are 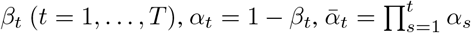. A neural network is then trained to reverse this process:

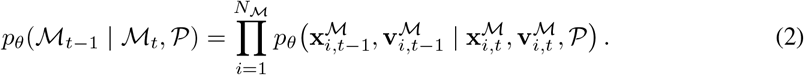

## 4 Methods

In this study, we first curate a large-scale, high-quality dataset linking protein-protein and protein-ligand complexes at shared orthosteric binding sites. Next, we introduce a diffusion-based model that conditions ligand generation on both pocket structure and peptide-defined PPI interactions (as shown in Figure 2), enabling joint pocket- and peptide-guided design of small-molecule PPI inhibitors.

**Figure 2:**
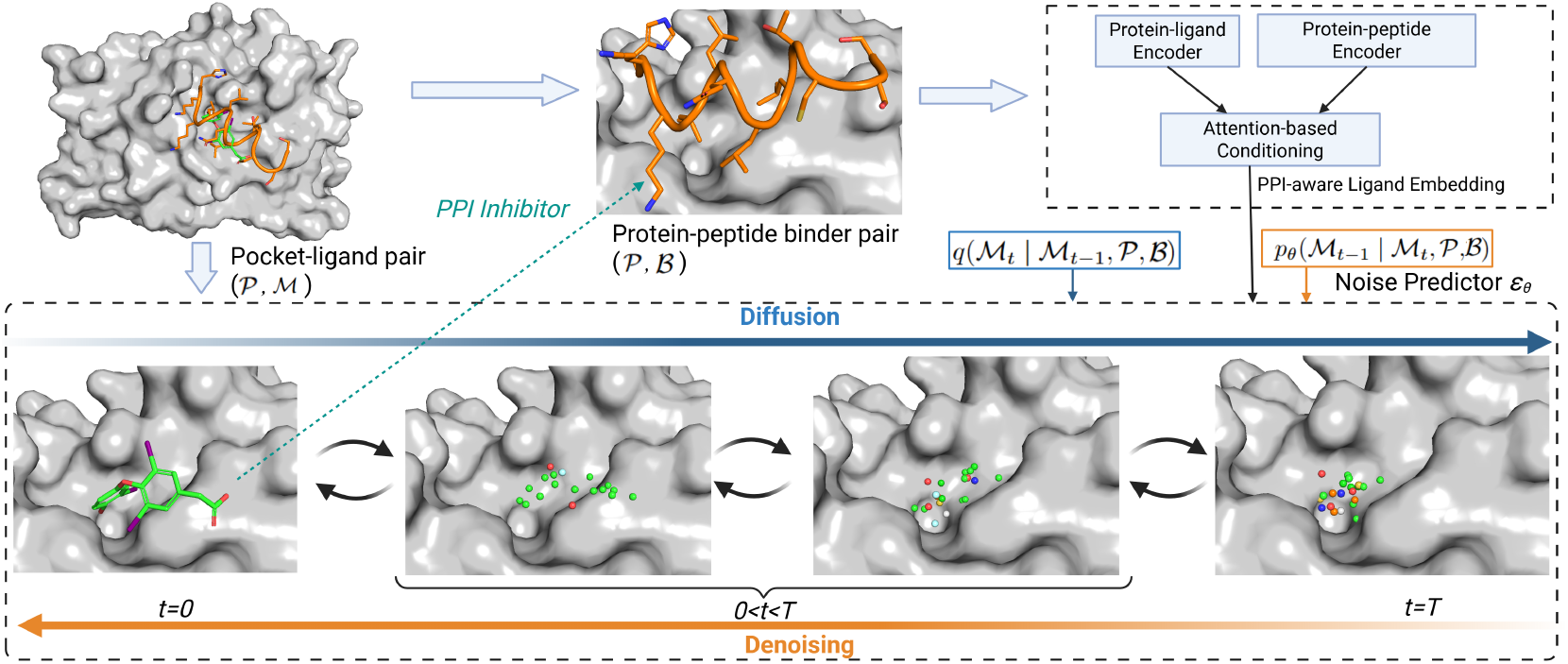
Schematic overview of Pep2Mol framework. PPI and PLI complexes are aligned based on the receptor protein binding regions. The diffusion process injects noise into the small-molecule PPI inhibitor. The generative process learns to denoise the corrupted inputs and recover the inhibitor atoms types and poses. PPI-derived interaction features are integrated with PLI representations via an attention mechanism, jointly guiding inhibitor generation throughout the diffusion process.

### 4.1 Data Preparation

#### Data Collection and Preprocessing

Our Pep2Mol dataset builds upon a recently published pocket-centric resource related to PPIs and PPI-related ligand binding sites Moine-Franel et al. [2024]. Based on the orthosteric annotations, the corresponding raw PDB structures of PLI and PPI pairs are retrieved from the RCSB Protein Data Bank, and each PPI–PLI pair is structurally aligned using the shared receptor as the reference. The ligand-binding pockets are defined as receptor residues located within 10 Å of the ligand. To enable joint pocket- and peptide binder-guided ligand generation, peptide segments are further extracted based on their proximity to the ligand-binding pocket, using a 10 Å interatomic distance cutoff, thereby establishing paired pocket–ligand–peptide correspondences for model training and inference. The resulting curated dataset contains 10,956 PPI-PLI pairs, comprising 688 unique PPI complexes, 1,654 unique PLI complexes, and 1,452 unique small molecules. The pipeline of data collection and pre-processing is detailed in Figure S1. To the best of our knowledge, this is the largest orthosteric PPI inhibitor dataset to date and provides paired pocket–ligand–peptide correspondences that directly enable structure-based generative modeling of orthosteric PPI inhibitors guided by peptide binders. Compared with the widely used SBDD benchmark CrossDocked2020, our PPI inhibitor-focused dataset exhibits a distinct pocket size–ligand length distribution (Figure S2): for comparable pocket sizes, ligands in Pep2Mol dataset tend to contain more heavy atoms, consistent with the shallow and extended geometry characteristic of PPI interfaces. To account for this difference, all benchmarked methods in this study sample the number of ligand heavy atoms from our dataset-specific ligand-size prior conditioned on pocket size for molecular generation.

#### Data Split

To establish a robust benchmark and minimize structural and chemical data leakage, binding pockets and ligands are first jointly clustered using hierarchical agglomerative clustering based on the TM-score with a cutoff 0.5 and Morgan-fingerprint Tanimoto similarity with a cutoff 0.7, respectively. This procedure ensures that structurally or chemically similar samples are assigned to the same cluster. Samples are considered eligible for the test set if their ligand satisfies the near drug-like criteria introduced by Chopra et al. [2025] and every residue in the PLI pocket has an identical amino-acid match in the PPI interface. The residue-level consistency minimizes ambiguity arising from potential mutations at the receptor-binding interfaces between the paired PLI and PPI structures. Eligible samples are then greedily assigned to the test set, prioritizing smaller clusters, until 100 PLI complexes are covered; the remaining samples from those clusters are excluded from the benchmark to prevent data leakage. The clusters not selected for the test set are subsequently partitioned into training and validation subsets. The final benchmark dataset contains 9,708 training, 1,079 validation, and 100 test samples.

### 4.2 Pep2Mol Framework

Pep2Mol is a diffusion-based generative framework for orthosteric PPI inhibitor design, where small-molecule ligands are generated under joint guidance from protein pocket structure and peptide-defined PPI interactions. The model follows a conditional DDPM formulation in which ligand atomic coordinates and atom types are iteratively denoised over diffusion timesteps. In this process, the ligand is treated as the only stochastic component, while the protein pocket and peptide binders remain fixed and provide structural conditioning. The architecture consists of three main components: (i) a PPI encoder, (ii) a PLI encoder, and (iii) a PPI-aware denoising network.

Following Guan et al. [2023a], each atom is represented by its 3D coordinates and feature embeddings. Ligand atoms use atom-type embeddings only, while pocket and peptide atoms use fused atom-type and residue-type embeddings to capture both atom- and residue-level context. Ligand atom types comprise one-hot vectors of chemical element type and aromatic information. Protein atom features include chemical element type, amino acid identity, and a binary indicator denoting whether the atom belongs to the protein backbone.

In contrast to conventional pocket-conditioned generation, Pep2Mol incorporates the peptide binder *B* as an additional condition, thereby leveraging peptide-specific interactions and binding pocket information to jointly guide small-molecule inhibitor design at PPI interfaces:

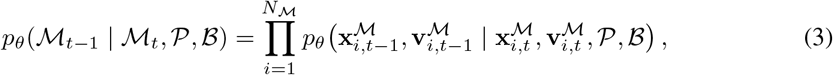

where *B* is integrated via the PPI encoder and attention-based conditioning described next. At each denoising step, Pep2Mol uses PPI-aware conditioning: the ligand branch is refined by attention under PPI guidance, while the pocket branch is updated before denoising. This joint conditioning enables the model to simultaneously incorporate orthosteric pocket geometry and peptide binding pattern, guiding ligand generation toward small-molecule inhibitors that competitively target PPI interfaces.

#### PPI Encoder

PPI encoder is implemented as an SE(3)-equivariant graph neural network operating on the PPI binding interface. Aligned pocket *P* and peptide *B* are taken as input. For pocket atoms, atom-type and residue-type embeddings are fused as 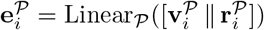 for feature concatenation. Similarly, for peptide atoms, atom-type and residue-type embeddings are fused as 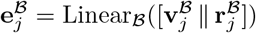. These embeddings are integrated into a unified PPI-conditioning graph with node features 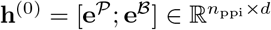, where *n*_ppi_ = *N*_*P*_ + *N*_*B*_, and *d* denotes the hidden feature dimension, and coordinates 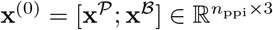. Edges defined by spatial proximity form the graph ℰ_ppi_. Positional relationships are encoded using L2 distances as scalar edge features for message passing, and displacement vectors (**x**_*j*_ −**x**_*i*_) for coordinate updates. These features are processed by SE(3)-equivariant message-passing blocks followed by graph attention layers:

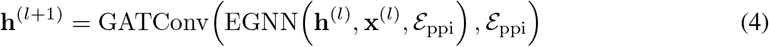

where 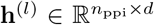and 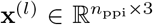 node features and atomic coordinates at layer *l*. After *L* stacked layers, the resulting features are used to form branch-specific PPI prompt representations 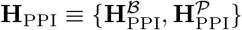, where 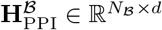 and 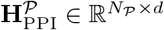 provide peptide-guiding and pocket-guiding conditioning signals in the diffusion model, respectively.

#### PLI Encoder

PLI encoder adopts a similar SE(3)-equivariant architecture and is responsible for encoding ligand atoms and orthosteric pocket atoms. The ligand atom-type embeddings are used directly, and the pocket atom-type and residue-type embeddings are fused as 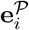. A binary identity embedding is concatenated to distinguish ligand and pocket atoms, followed by a linear projection to obtain initial node features. Ligand and pocket atoms are combined into a single complex graph with edges defined by spatial proximity. Relative coordinate differences and distances are computed and passed to SE(3)-equivariant message-passing blocks, which update both node features and atomic coordinates. After equivariant updates, geometric attention is applied using graph attention layers to enhance local spatial features. The resulting ligand and pocket representations are further fused through an additional attention-based graph layer to model cross-interface interactions. The encoder outputs separate representations for ligand and pocket atoms: 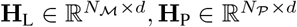, which together define the PLI conditioning representation **H**_PLI_ *≡{***H**_L_, **H**_P_*}*.

#### Encoder Conditioning in Diffusion

At diffusion step *t*, the denoising network receives noisy ligand states together with conditioning features from the PLI and PPI encoders. The PLI encoder provides ligand-side and pocket-side features (**H**_L_, **H**_P_), while the PPI encoder provides peptide-guiding and pocket-guiding prompt features 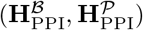. The perturbed ligand atom types 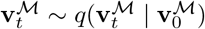 and diffusion step *t* are projected to form 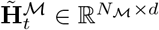; pocket atom types **v**^*P*^ and residue features **r**^*P*^ are projected to form 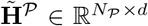. The ligand branch is updated by attention-based fusion, whereas the pocket branch is updated by projection-based fusion:

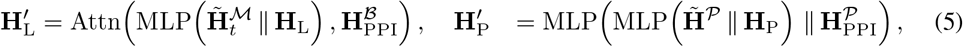

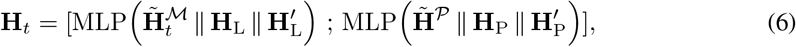

where 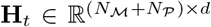 denotes the fused PLI–PPI conditioning representation passed to the denoising network. Formally, the denoising model predicts:

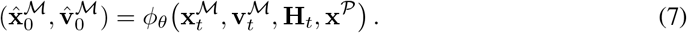

For PLI and PPI encoders, SE(3)-equivariance is maintained by two separate EGNNs. The GATConv layer operates only on the SE(3)-invariant hidden states, rather than directly updating geometric features such as coordinates, thereby preserving SE(3)-equivariance. Similarly, the attention-based fusion uses only SE(3)-invariant features output by the PPI encoder, which are coordinate-system agnostic, and integrates them with the invariant features from the PLI encoder through cross-attention. Therefore, relative positional information of PPI is removed, while only interaction information encoded in feature space is retained and fused with the PLI representation as joint conditioning. The full training procedure is summarized in Algorithm S1 in Supplementary materials.

### 4.3 Training Objective

Ligand coordinates and atom types are noised at timestep *t* during training. The model predicts denoised ligand coordinates and clean atom-type logits in Eqn. (7). The atom-type loss trains the model-induced reverse categorical distribution to match the posterior 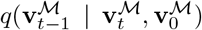. The training loss combines a coordinate denoising loss and a categorical atom-type loss:

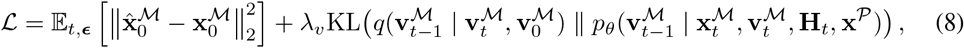

where *λ*_*v*_ balances the accuracy of atom positions and types.

## 5 Experiments

### 5.1 Experimental Settings

#### Datasets and Baseline Methods

Experiments are conducted on our curated dataset using the leakage-proof splitting strategy described in Sections 4.1. We compare our model with recent representative methods for SBDD, including diffusion-based methods TargetDiff Guan et al. [2023a], Pocket2Mol Peng et al. [2022], and IPDiff Huang et al. [2024b], as well as the flow matching-based framework DrugFlow Shen et al. [2024]. All baseline methods are evaluated under both training-from-scratch and fine-tuning settings. In the training-from-scratch setting, models are randomly initialized and trained directly on our dataset. In the fine-tuning setting, baseline models are initialized from their official checkpoints pretrained on CrossDocked2020 dataset and subsequently fine-tuned on our Pep2Mol benchmark. One concurrent work, Peptide2Mol He et al. [2025] is conceptually related in leveraging peptide binders for ligand generation, but is excluded from comparison due to its preprocessing incompatible with our competitive binding setup. Its requirement to merge protein, peptide, and ligand into a single SDF file introduces severe atomic clashes on our dataset, where ligands and peptides competitively bind the same orthosteric interface, leading to widespread sample failures during training and inference.

#### Evaluation Metrics

The generated ligands are comprehensively evaluated from three perspectives: target binding affinity, molecular properties, and molecular structures. Target binding affinity is assessed for each valid ligand–protein pair using AutoDock Vina Eberhardt et al. [2021] under three settings: fixed-pose scoring (Vina Score), local pose optimization (Vina Min), and global docking optimization (Vina Dock). We also report a High-Affinity metric defined as the proportion of generated ligands with Vina Scores lower than those of the reference ligands in the same binding pocket. Molecular properties are evaluated using QED Bickerton et al. [2012] for drug-likeness and SA scores Ertl and Schuffenhauer [2009] for synthetic accessibility. Molecular structure is assessed by computing Earth Mover’s Distances (EMDs) between the all-atom pairwise distance distributions of generated and reference ligands, following Vignac et al. [2023] and O Pinheiro et al. [2023], with all atom pairs considered jointly without bond-type weighting.

### 5.2 Main Results

#### Target Binding Affinity and Molecule Properties

We first compare Pep2Mol with baseline models using metrics related to binding affinity and molecular properties. As shown in Table 1, Pep2Mol achieves the strongest binding performance among all evaluated methods. It obtains average Vina Score, Vina Min, and Vina Dock values of −6.93, −8.03, and −9.03, respectively, outperforming the second-best results: −4.75, −6.36, −7.45 from DrugFlow. Pep2Mol also reaches the highest High-Affinity ratio of 0.76 *±*0.05 (mean *±* std), representing a substantial improvement over the best-performing baseline, IPDiff, which obtains 0.50 *±* 0.02. Median docking scores further confirm this trend, with Pep2Mol achieving the best Vina Score, Vina Min, and Vina Dock values of −7.81, −8.29, and −8.95 respectively, as shown in Table S1. The significant affinity improvements achieved by Pep2Mol indicate that peptide-derived structural guidance effectively enhances PPI-targeted ligand generation and improves the predicted binding affinity of the designed ligands. In contrast, conventional pocket-conditioned SBDD models that omit peptide context generally underperform both reference ligands and Pep2Mol-designed ligands, highlighting their limited effectiveness for the large, shallow, and chemically complex PPI interface.

**Table 1:** Comparison of ligand generation performance. Baselines marked with * are fine-tuned from official pretrained checkpoints, while unmarked models are trained from scratch on our dataset. Results are reported as mean (std) over the 3 runs per model. The best value in each column is highlighted in bold, and the second-best value is underlined.

| Model | Vina Score (↓) | Vina Min (↓) | Vina Dock (↓) | QED (↑) | SA (↑) | High-Affi. (↑) |
| --- | --- | --- | --- | --- | --- | --- |
| Reference | -4.99 | -6.36 | -7.14 | 0.67 | 0.83 | – |
| Pocket2Mol | -3.24 (0.18) | -4.79 (0.14) | -5.52 (0.31) | <u>0.55 (0.01)</u> | <u>0.83 (0.01)</u> | 0.20 (0.02) |
| Pocket2Mol* | -3.95 (0.07) | -5.32 (0.05) | -6.03 (0.17) | **0.58 (0.01)** | **0.87 (0.01)** | 0.26 (0.02) |
| TargetDiff | -2.56 (0.31) | -5.06 (0.09) | -6.92 (0.06) | 0.49 (0.01) | 0.60 (0.01) | 0.43 (0.03) |
| TargetDiff* | -4.38 (0.27) | -5.55 (0.21) | -6.88 (0.15) | 0.39 (0.01) | 0.54 (0.02) | 0.46 (0.05) |
| IPDiff | -4.17 (0.68) | -5.18 (0.30) | -6.59 (0.16) | 0.37 (0.02) | 0.54 (0.02) | 0.49 (0.01) |
| IPDiff* | -4.35 (0.34) | -5.58 (0.04) | -6.89 (0.18) | 0.41 (0.06) | 0.58 (0.01) | 0.50 (0.02) |
| DrugFlow | -2.84 (0.08) | -5.54 (0.29) | -7.02 (0.42) | 0.45 (0.06) | 0.56 (0.00) | 0.27 (0.02) |
| DrugFlow* | -4.75 (0.26) | -6.36 (0.26) | -7.45 (0.25) | 0.51 (0.04) | 0.63 (0.01) | 0.41 (0.03) |
| Pep2Mol | **-6.93 (0.30)** | **-8.03 (0.74)** | **-9.03 (0.67)** | 0.38 (0.02) | 0.52 (0.02) | **0.76 (0.05)** |

Regarding molecular properties, Pep2Mol-generated molecules exhibit QED and SA values comparable to those of other diffusion-based baselines, including IPDiff and TargetDiff, although they lag behind the autoregressive model Pocket2Mol and the flow-matching-based model DrugFlow. Consistent with previous studies Huang et al. [2024a], Guan et al. [2023b], we observe a trade-off between binding-related and property-related metrics. Accordingly, we place less emphasis on QED and SA, as these metrics are typically used as approximate screening indicators in practical drug discovery and are generally acceptable when maintained within a reasonable range.

#### Molecular Structure

We evaluate Pep2Mol against baseline methods in terms of structural plausibility of the generated molecules. Figure 3 presents the empirical distributions of all-atom pairwise distances for the generated molecules and compares them with those of the reference ligands. The Earth Mover’s Distance (EMD) between the generated and reference distributions is computed to quantify the minimum transport cost between them while providing a continuous measure under distributional sparsity. As shown in Figure 3, Pep2Mol achieves the lowest EMD of 0.33 relative to the ground-truth distribution, outperforming the next-best baseline method, DrugFlow (0.43), and indicating its strong ability to capture realistic atomic distance patterns. Results are obtained across the three independent runs per model.

**Figure 3:**
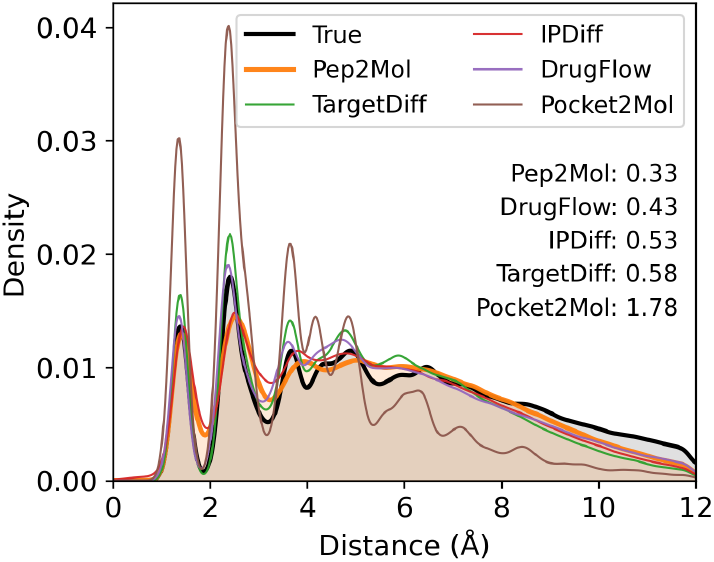
All-atom pairwise distance distributions for reference molecules in the test set (black) and molecules generated by different methods. Distributional similarity is quantified with EMD.

#### Case Studies

Figure 4 presents representative case studies, comparing Pep2Mol with the strong diffusion-based baseline IPDiff. The interaction similarity of the generated ligands to the ground truth peptides is quantified by the Jaccard similarity of the contact residues identified using Protein-Ligand Interaction Profiler Schake et al. [2025] with 4 Å as cutoff. Higher scores indicate better preservation of peptide-guided interactions. The ground truth columns show how the reference ligands and peptides define and compete for the orthosteric protein binding interface. Compared with the baseline method, Pep2Mol-generated ligands more consistently preserve peptide-engaged interaction residues and exhibit higher interaction similarity across all samples. These results further demonstrate that peptide binder conditioning enables Pep2Mol to capture peptide binding patterns for inhibitor design at the PPI interfaces.

**Figure 4:**
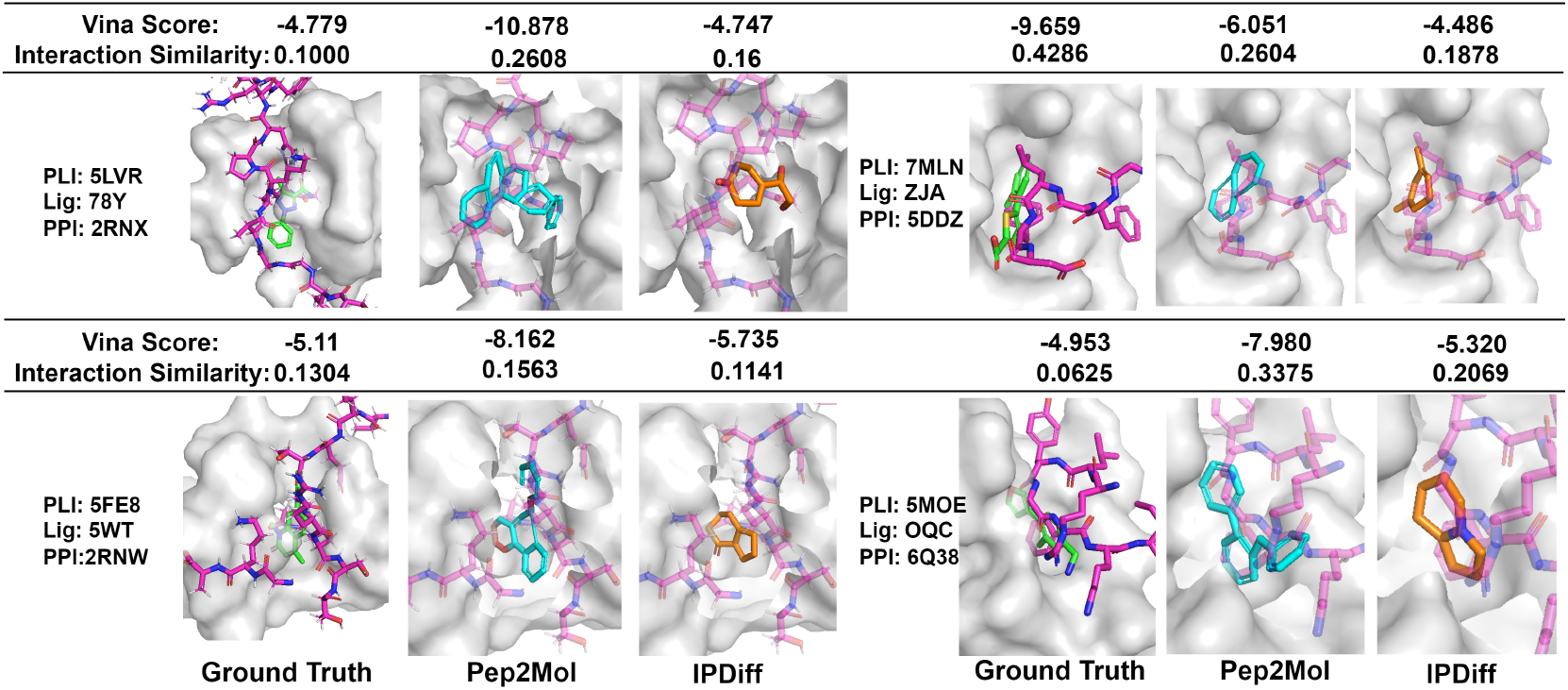
Case studies of peptide-guided ligand generation. Examples show protein-peptide interfaces alongside ground-truth pocket-ligand complexes and generated ligands. Interaction similarity highlights how peptide binders guide generated ligands to preserve important interaction residues.

#### Ablation Study

We further conduct an ablation study to evaluate the contribution of peptide-derived conditioning. Specifically, we compare Pep2Mol with a variant (*w/o PPI*), where both peptide inputs and the PPI encoder are removed, leaving the model conditioned only on the receptor pocket, as in conventional SBDD baselines. As shown in Table 2, this variant consistently underperforms

**Table 2:** Ablation study on peptide conditioning using mean values.

| Model | Vina Score ↓ | Vina Min ↓ | Vina Dock ↓ | QED ↑ | SA ↑ | High-Affi. ↑ |
| --- | --- | --- | --- | --- | --- | --- |
| Pep2Mol | **-6.93(0.30)** | **-8.03(0.74)** | **-9.03(0.67)** | 0.38(0.02) | 0.52(0.02) | **0.76(0.05)** |
| w/o PPI | -4.99(0.42) | -5.51(0.47) | -6.72(0.45) | **0.40(0.11)** | **0.57(0.04)** | 0.54(0.02) |

Pep2Mol across target binding affinity-related metrics and achieves performance comparable to other SBDD baselines. The average Vina Score, Vina Min, and Vina Dock values decrease markedly from −4.99 to −6.93, from −5.51 to −8.03, and from −6.72 to −9.03, respectively. The same trend is also observed for median docking scores (Table S2), where Pep2Mol achieves median Vina Score, Vina Min, and Vina Dock of −7.81, −8.29, and −8.95, outperforming the *w/o PPI* variant at −5.43, −5.70, and −6.71. This ablation study demonstrates that binding-peptide conditioning enables Pep2Mol to generate PPI-targeting small molecules with higher predicted binding affinity, highlighting the key contribution of the peptide-derived structural guidance.

## 6 Conclusion and Limitation

In this study, we present Pep2Mol, a full-atom SE(3)-equivariant diffusion framework for three-dimensional molecule design that targets orthosteric PPI binding sites by explicitly incorporating peptide or protein binders as structural guidance. We also establish the largest high-quality, machine learning-ready dataset to date of experimentally resolved orthosteric PPI-PLI pairs, providing a foundational resource for structure-based PPI inhibitor design and opening new avenues for AI-driven drug discovery. Extensive evaluations demonstrate that Pep2Mol outperforms existing state-of-the-art methods, generating chemically valid PPI inhibitors with improved predicted binding affinity. Despite these advances, Pep2Mol has several limitations. First, it is more computationally demanding during both training and sampling than conventional pocket-conditioned methods, due to the additional peptide-conditioned atomic inputs and the corresponding PPI encoder. Future work will improve scalability through more efficient graph construction strategies, such as atom-block hierarchical representations. Second, Pep2Mol is an SBDD method that requires a peptide binder structure as input, which may limit direct application to targets without resolved complex structures. When only the receptor structure is available, the complex structure could first be obtained using co-folding methods or protein docking, after which Pep2Mol can be applied. Finally, from a molecular-generation perspective, incorporating bond-aware diffusion and connectivity constraints represents a promising direction to further improve drug-likeness and energetic realism.

## 7 Acknowledgements

This work was supported in part by the University of Florida (UF Startup Fund, UF Health Cancer Institute Pilot Grant #CAT-2026-20, and UF-NVIDIA Artificial Intelligence and Complex Computational Research Award to Y. L.), National Institutes of Health (R21EB037868 to Y. L.), and the Nicholas Bodor Professorship Fund (to C. L.). Research reported in this publication was supported in part by the UF Health Cancer Institute, supported in part by state appropriations provided in Fla. Stat. § 381.915 and the National Cancer Institute of the National Institutes of Health under Award Number P30CA247796. The content is solely the responsibility of the authors and does not necessarily represent the official views of the National Institutes of Health or the State of Florida.

## A Supplementary materials

### A.1 Curation of Pep2Mol dataset

The detailed procedures for curating the Pep2Mol dataset are shown in Figure S1.

**Figure S1:**
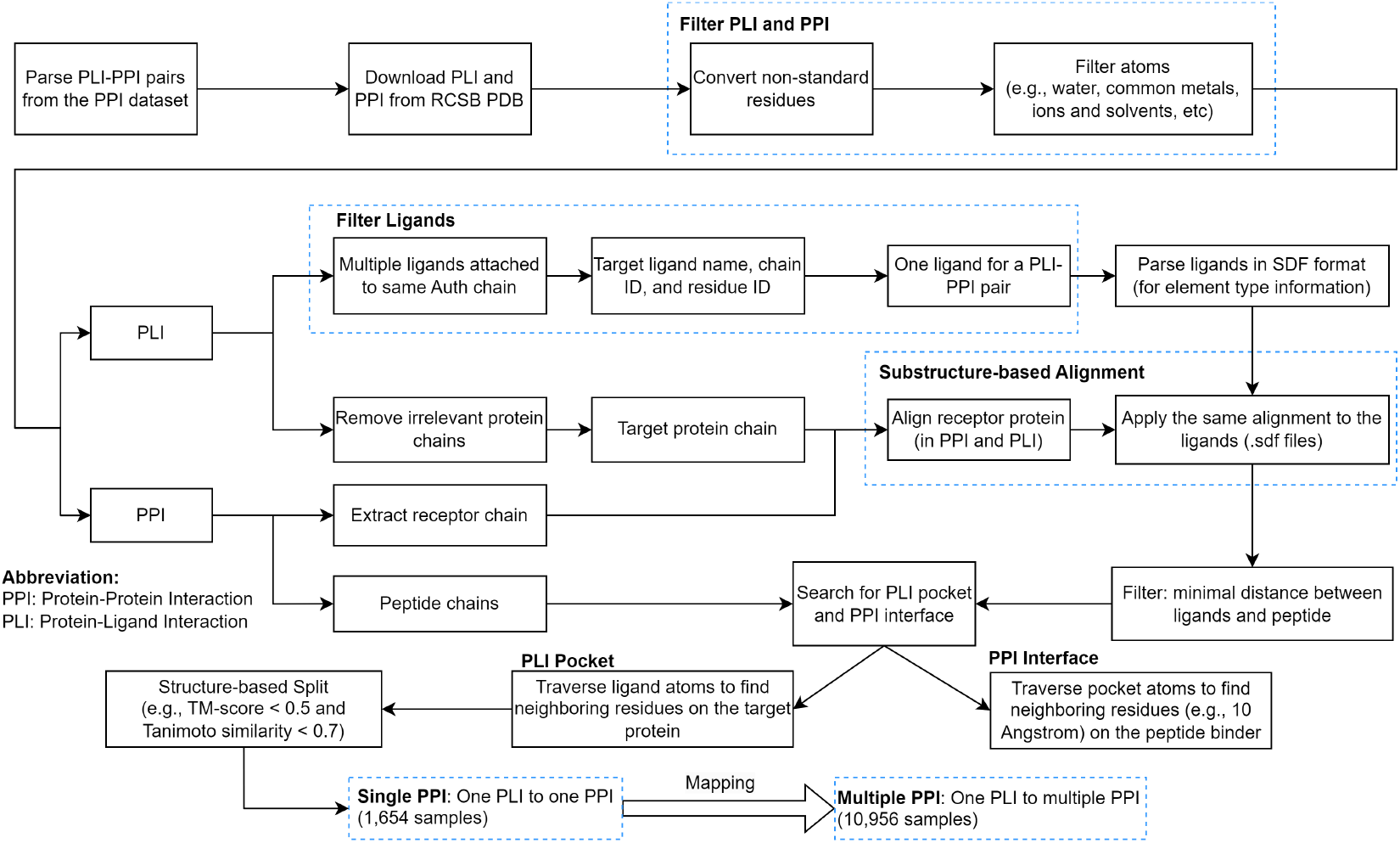
The flowchart of data collection and preprocessing for the Pep2Mol dataset.

### A.2 Comparison between Pep2Mol and CrossDocked2020 datasets

We compare the relationship of pocket size and ligand size in Pep2Mol and CrossDocked2020 datasets. Pocket size is defined as the spatial extent of the ligand-binding pocket and is computed, following Guan et al. [2023a], as the median of the largest pairwise distances among pocket atoms. Ligand size is defined as the number of heavy atoms in the small-molecule ligand. Figure S2 compares the pocket-size-conditioned ligand-size prior fitted on the Pep2Mol dataset with that fitted on CrossDocked2020, showing that, for pockets of comparable size, the Pep2Mol-specific prior tends to sample larger ligands.

**Figure S2:**
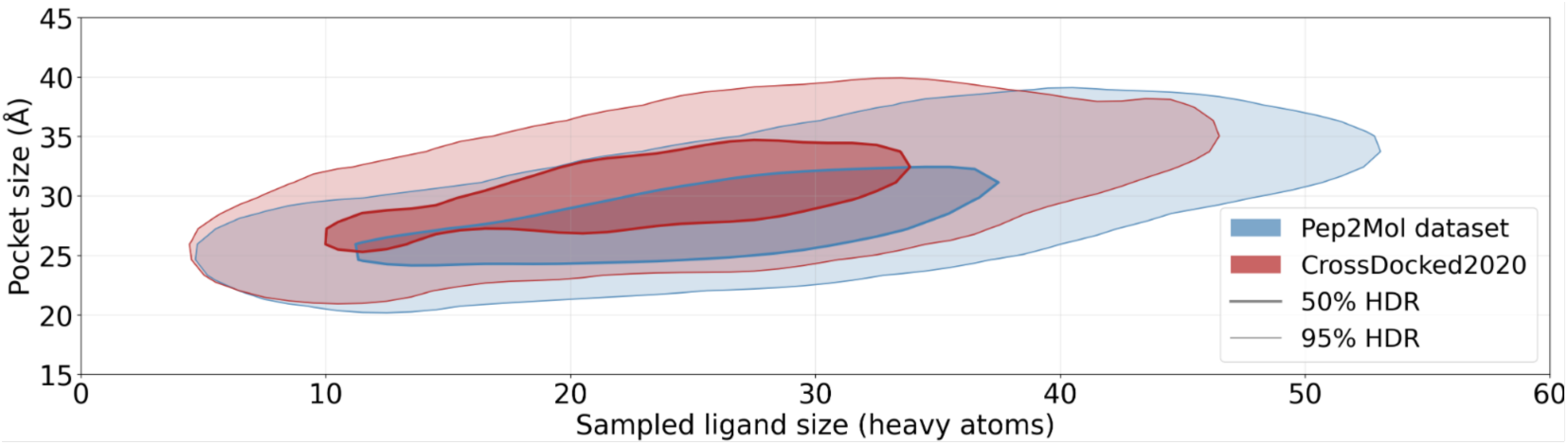
Sampled ligand-size vs. pocket size under dataset-specific priors. Ligand sizes sampled from each dataset’s fitted pocket-size-conditioned prior; filled regions and contours mark the 50% and 95% highest-density regions (HDR) for Pep2Mol (blue) and CrossDocked2020 (red), e.g., the 50% HDR encloses half of each prior’s samples.

### A.3 Training process

#### Algorithm S1

Training Procedure of PPI-Aware Pep2Mol

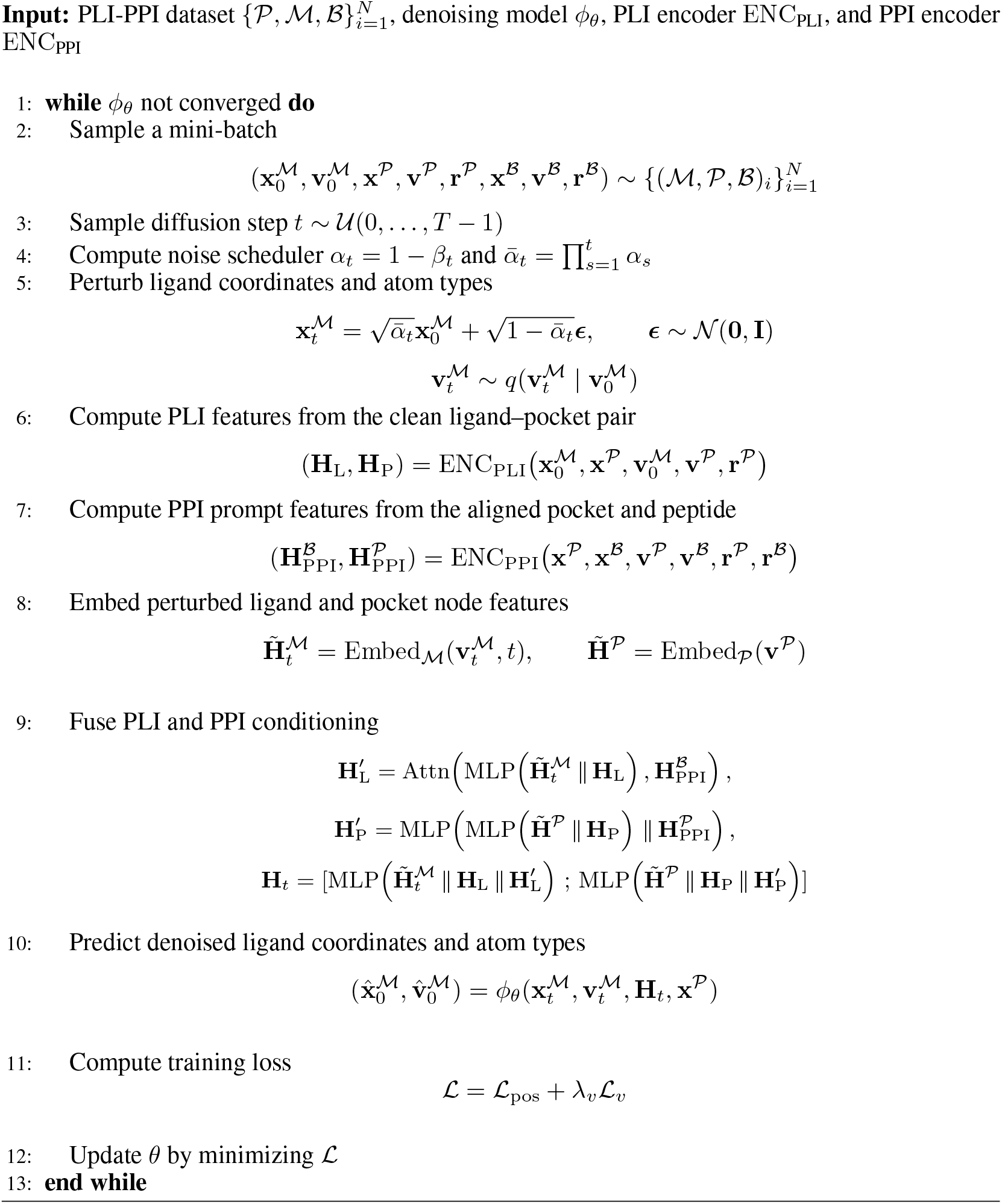

where the variables are:

- 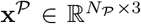: 3D coordinates of atoms in the aligned receptor pocket, also used as the peptide-interface representation.
- 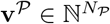 : atom-type indices of pocket atoms.
- 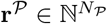 : residue-type indices of pocket atoms.
- 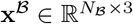: 3D coordinates of atoms in the aligned peptide.
- 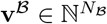 : atom-type indices of peptide atoms.
- 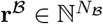 : residue-type indices of peptide atoms.
- 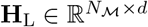: ligand-side conditioning features produced by the PLI encoder
- 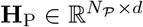: pocket-side conditioning features produced by the PLI encoder
- 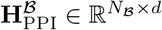: peptide-guiding prompt features used in ligand-side fusion
- 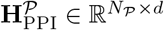: pocket-guiding prompt features used in protein-side fusion
- 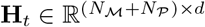: final fused PLI–PPI conditioning representation passed to the denoising network.

### A.4 Comparison of median Vina scores

Comparison of ligand generation performance using median docking scores is shown in Table S1. More setting details can be found around Table 1 in the main text. For the ablation study in Table 2, its median values are reported in Table S2.

**Table S1:** Comparison of ligand generation performance using median docking scores. Results are reported as median (std) over the 3 runs per model.

| Model | Vina Score (↓) | Vina Min (↓) | Vina Dock (↓) | QED (↑) | SA (↑) | High-Affi. (↑) |
| --- | --- | --- | --- | --- | --- | --- |
| Reference | -5.20 | -6.15 | -6.95 | 0.70 | 0.84 | – |
| Pocket2Mol | -3.34 (0.06) | -4.68 (0.14) | -5.52 (0.17) | 0.53 (0.03) | 0.83 (0.01) | 0.20 (0.02) |
| Pocket2Mol* | -4.08 (0.04) | -5.33 (0.08) | -6.05 (0.09) | **0.57 (0.01)** | **0.89 (0.01)** | 0.26 (0.01) |
| TargetDiff | -4.88 (0.15) | -5.83 (0.16) | -7.02 (0.06) | 0.50 (0.01) | 0.59 (0.01) | 0.42 (0.04) |
| TargetDiff* | -5.13 (0.22) | -5.68 (0.11) | -6.90 (0.12) | 0.39 (0.01) | 0.54 (0.02) | 0.44 (0.02) |
| IPDiff | -5.07 (0.54) | -5.19 (0.49) | -6.57 (0.41) | 0.35 (0.03) | 0.56 (0.03) | 0.45 (0.04) |
| IPDiff* | -5.25 (0.34) | -5.82 (0.23) | -6.92 (0.23) | 0.41 (0.10) | 0.57 (0.01) | 0.50 (0.02) |
| DrugFlow | -3.70 (0.09) | -5.60 (0.33) | -7.15 (0.41) | 0.43 (0.07) | 0.55 (0.01) | 0.27 (0.02) |
| DrugFlow* | -4.95 (0.22) | -6.31 (0.20) | -7.40 (0.22) | 0.51 (0.06) | 0.63 (0.01) | 0.41 (0.03) |
| Pep2Mol | **-7.81 (0.52)** | **-8.29 (0.80)** | **-8.95 (0.81)** | 0.38 (0.00) | 0.51 (0.03) | **0.76 (0.04)** |

**Table S2:**
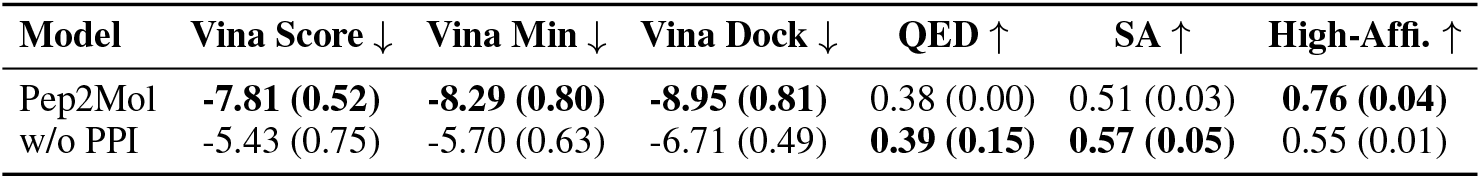
Ablation study on peptide conditioning using median values.

| Model | Vina Score ↓ | Vina Min ↓ | Vina Dock ↓ | QED ↑ | SA ↑ | High-Affi. ↑ |
| --- | --- | --- | --- | --- | --- | --- |
| Pep2Mol | **-7.81 (0.52)** | **-8.29 (0.80)** | **-8.95 (0.81)** | 0.38 (0.00) | 0.51 (0.03) | **0.76 (0.04)** |
| w/o PPI | -5.43 (0.75) | -5.70 (0.63) | -6.71 (0.49) | **0.39 (0.15)** | **0.57 (0.05)** | 0.55 (0.01) |

### A.5 Near drug-like property filter

Near drug-like properties are listed in Table S3 according to Chopra et al. [2025]. Compare to “Drug-like” and “Lead-like”, these are more relaxed filters to classify ligands.

**Table S3:**
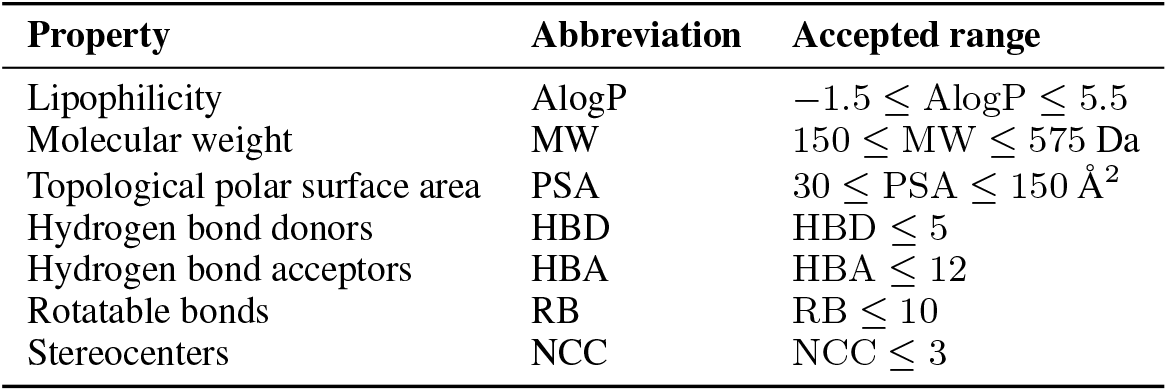
Near drug-like filtering criteria used for test-set ligand selection.

| Property | Abbreviation | Accepted range |
| --- | --- | --- |
| Lipophilicity | AlogP | $-1.5 \leq \text{AlogP} \leq 5.5$ |
| Molecular weight | MW | $150 \leq \text{MW} \leq 575 \text{ Da}$ |
| Topological polar surface area | PSA | $30 \leq \text{PSA} \leq 150 \text{ \AA}^2$ |
| Hydrogen bond donors | HBD | $\text{HBD} \leq 5$ |
| Hydrogen bond acceptors | HBA | $\text{HBA} \leq 12$ |
| Rotatable bonds | RB | $\text{RB} \leq 10$ |
| Stereocenters | NCC | $\text{NCC} \leq 3$ |

**Table S4:** Efficiency and model-size comparison.

| Model | Train Time | Sample / Attempt | Params |
| --- | --- | --- | --- |
| Pocket2Mol | 1.3h | 3s | 3.7M |
| TargetDiff | 6h | 12s | 2.8M |
| IPDiff | 6.2h | 15s | 3.5M |
| DrugFlow | 3.5h | 12s | 12.1M |
| Pep2Mol | 25h | 51s | 4.3M |

### A.6 Computational resource

Pep2Mol is trained on two NVIDIA B200 GPUs using the Adam optimizer with an initial learning rate of 5 *×*10^−4^. A plateau-based learning rate scheduler is applied for decay, with a minimum learning rate of 1*×* 10^−6^. Training is performed with a batch size of 1 and typically converges within 600 epochs. The efficiency and model-size comparison is reported in Table S4.

### A.7 Hyperparameter settings

Hyperparameter settings used in Pep2Mol is listed in Table S5.

**Table S5:** Hyperparameters for Pep2Mol.

| Hyperparameter | Value | Description |
| --- | --- | --- |
| hidden_dim | 128 | hidden dimension in Pep2Mol |
| n_layers | 9 | number of GNN layers in Pep2Mol |
| k_neighbors | 32 | number of neighbors per node in the KNN graph |
| beta_start | $1 \times 10^{-7}$ | starting $\beta$ value in the diffusion schedule |
| beta_end | $2 \times 10^{-3}$ | ending $\beta$ value in the diffusion schedule |
| sampling_steps | 1000 | number of diffusion sampling steps |
| atom_type_loss_weight | 100 | weight of the atom-type prediction loss term |

